# Elucidating the Assembly of Gas Vesicles by Systematic Protein-Protein Interaction Analysis

**DOI:** 10.1101/2023.07.23.550228

**Authors:** Manuel Iburg, Andrew P. Anderson, Vivian T. Wong, Erica D. Anton, Art He, George J. Lu

## Abstract

Gas vesicles (GVs) are gas-filled microbial organelles formed by unique 3-nm thick, amphipathic, force-bearing protein shells, which can withstand multiple atmospheric pressures and maintain a physically stable air bubble with megapascal surface tension. However, the molecular process to assemble this shell remains elusive: while 6-8 assembly factor proteins were identified as essential, none of them have a defined function. As one of the first steps to elucidate the assembly mechanism, we devise a high-throughput *in vivo* assay to determine the interactions of all 11 proteins in a GV operon. Complete or partial deletions of the operon establish the interdependence relationship of the interaction on the background GV proteins with additional information on assembly tolerance and cellular burden. Clusters of GV protein interactions are revealed, which establishes the plausible protein complexes important for the assembly process of these protein organelles. We anticipate our findings will set the stage for solving the molecular mechanism of GV assembly and designing GVs that efficiently assemble in heterologous hosts during biomedical applications.

## INTRODUCTION

Gas vesicles (GVs) are a class of gas-filled protein organelles evolved in photosynthetic microbes, which use them as flotation devices to compete for the surface of the water to maximize photosynthesis^2,3^. Besides this native function, the interest in GVs has grown substantially due to the recent development of a wide range of biomedical applications based on these genetically encodable nanostructures, including gene expression imaging by ultrasound, MRI, and optical methods^5-12^, protease sensing^14^, payload delivery^15^, cellular manipulation^16,17^, gas delivery^18,19^, cell tracking^20^, and pressure sensing^21^. This rapidly growing set of applications has now demanded a deeper understanding of the biology of GV formation to facilitate their molecular engineering.

As a product of evolution and natural selection, GVs have a fundamentally different design compared to man-made synthetic bubbles. Bubbles usually trap air inside at a non-equilibrium state, leading to a finite lifetime, and moreover, as the diameter of bubbles reduces, a higher Laplace repressure builds up inside, leading to a faster dissipation of entrapped air^22^. For GVs, however, the protein shell is permeable to individual molecules of both gas and water, and GVs maintain an inner gas compartment by having a hydrophobic inner surface that prevents heterogeneous condensation of water molecules into liquid. Meanwhile, the small size of the gas compartments minimizes the chance of homogeneous condensation^23^. In combination with a rigid shell that prevents the shrinking of the gas compartments, GVs outcompete the best synthetic bubbles in both the lifetime and the size – nanobubbles have only recently reached < 200 nm in diameter and a lifetime of minutes^24^, while GVs can be made with diameters smaller than 100 nm and stable for months^25,26^.

This unique design principle of GVs necessitates a set of complex cellular machinery to support their assembly. For example, the major shell proteins have highly hydrophobic inner surfaces and aggregate even in the presence of strong detergents such as sodium dodecyl sulfate^27^. It is thus hypothesized that the shell proteins are coordinated constantly by other proteins from their synthesis at the ribosomes until their insertion into the GV nanostructures. Additionally, different from synthetic bubbles to which gas is exogenously provided, the gas inside GVs is equilibrated from that dissolved in the surrounding fluid, which hints at an interesting initiation process and the subsequent elongation of the gas compartment. Notably, this is in contrast to most of the other protein nanostructures described to date, which are formed by shell proteins with a propensity of self-assembly^28-32^.

Despite the recognition of the unique assembly mechanism, the understanding of the functional roles of the assembly factor proteins and the molecular assembly process has severely lagged behind. Only the major shell protein, which is usually denoted as *gvpA* or *gvpB*, and the minor shell protein, *gvpC*, have clearly defined functions^1,33^. The remaining genes often have undefined functions in a given GV operon, which usually contains on the order of 10 genes^2,12^. Taking one of the most well-studied GV operons, the 9-gene *gvpACNJKGFVW* from *Anabaena flos-aquae* as an example, the 7 genes after *gvpA* and *gvpC* were believed to be assembly factors or minor constituents of GVs. Among them, GvpF is the only one with high-resolution crystal structures determined from orthologs, but neither structure provided an explanation of how the protein may function in assembling GVs^34,35^. Much less is known for the other assembly proteins, and it is also puzzling to see the existence of this 7-gene cassette is contradicting the minimal 8 essential genes determined in the case of the pNL29 GV operon originally cloned from *Bacillus megaterium*^6^.

In this study, we set out to make one of the first steps toward understanding the GV assembly mechanism by establishing a systematic protein-protein interaction roadmap among all the GV proteins. In particular, considering the large number of assembly factor proteins needed to probe and that they may form multiprotein complexes with intricate interdependence relationships, we hypothesize that a unique strategy needs to be devised to allow a high-throughput genetic design, a workflow to assess multiple parameters, and the flexibility to probe the interdependence of protein-protein interactions. After establishing this workflow, the selective supplementation of the background GV proteins will allow us to tease out the interdependence relationship and establish subnetworks of interactions, leading to the formulation of hypotheses on proteins complexes responsible for key assembly stages, such as the initiation, elongation, chaperoning, and high-order packing. Together, these insights will lay the foundation for future studies of individual protein complexes and the establishment of a molecular mechanism of GV assembly.

## RESULTS

### Designing an efficient strategy to achieve full coverage of interaction conditions

Before designing an experimental strategy, the first decision to make is which GV genotype to focus on, and to this end, we chose the pNL29 operon originally cloned from *B. megaterium*^36^ (**Figures 1A-C**). Three reasons underlined this choice: (i) many recent biomedical applications including both bacterial and mammalian acoustic reporter genes^5,6^ use the assembly factor proteins of pNL29 operon, and thus the knowledge of these proteins will directly contribute to the engineering and design of GVs; (ii) the high-resolution structure of the GVs encoded by this operon was recently determined^1^, opening the door to the potential structure-based modeling of the assembly process using these structures; and (iii) pNL29 operon allows GVs to be heterologously expressed and assembled in *E. coli*^37^, which provides a robust platform to leverage genetic tools to study the assembly mechanism.

**Fig. 1.**
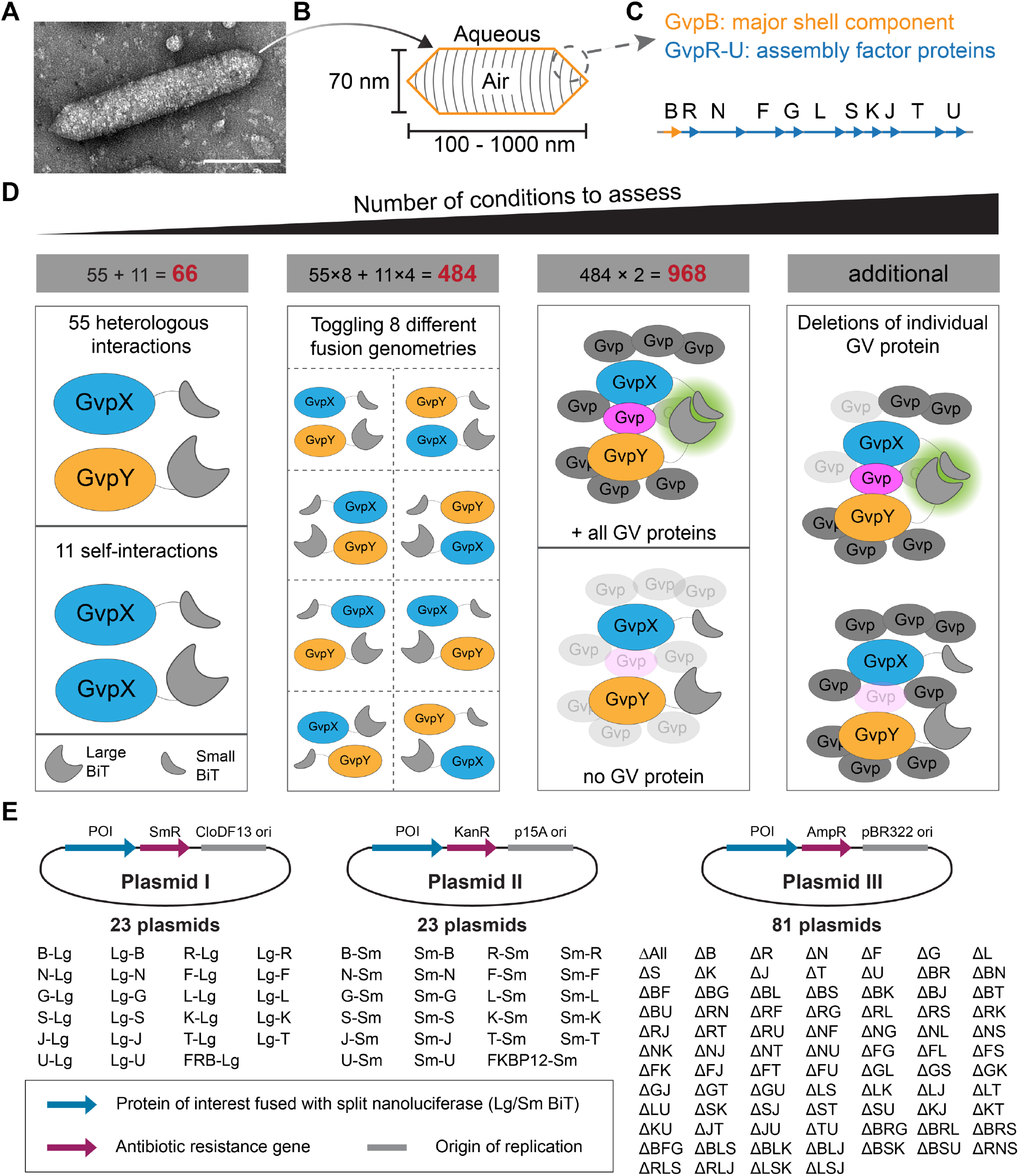
Designing an efficient strategy to achieve full coverage of interaction conditions. (**A** and **B**) A transmission electron microscopy (TEM) image and a schematic representation of GVs encoded by the pNL29 operon. (**C**) The operon consists of the structural component (labeled in yellow) and assembly factors (labeled in blue). (**D**) Calculating the numbers of genetic conditions through the 4 layers of variables. GV proteins fused to Large BiT and Small BiT of the split NanoLuc are marked in yellow and blue, respectively. In the 1^st^ layer, the heterologous and self-interactions of all 11 GV proteins result in 66 conditions, and in the 2^nd^ layer, toggling the fusion geometries brings the number to a total of 484 conditions excluding controls. In the 3^rd^ layer, to determine if protein-protein interaction is direct or interdependent on a third GV protein (marked in magenta), all interactions will be measured in the presence and absence of the other GV proteins (knocked-out GV proteins marked in translucent gray). If the interaction is dependent on a third protein, the bioluminescence signal (marked in green) will only show up when all GV proteins are present. In the 4^th^ layer, to search for the identity of the third GV protein (marked in magenta), partial knockout of GV proteins will be employed, which further expands the number of conditions. (**E**) Minimizing the number of plasmids needed to construct by the strategy of mix- and-match of 3 plasmids. The origins of replication, resistance markers, and proteins of interest (POI) are marked in gray, magenta, and blue, respectively. The abbreviated names of all 127 plasmids are listed. Lg: Large BiT. Sm: Small BiT. B, R, N, F, G, L, S, K, J, T, U: individual GV protein, for example, B stands for GvpB. Δ stands for a plasmid that contains all 11 GV proteins except for those indicated. See also the plasmid map in **Figure S1** and the full plasmid list in **Table S1**.

Following this choice, we conducted a back-of-the-envelope calculation on the number of conditions we would need to probe (**Figure 1D**). pNL29 operon contains 11 genes, and thus there is a total of 55 pairs of heterologous interactions. Additionally, we reasoned that some of the assembly factor proteins might form homo-oligomers that are important to assay, which added another 11 self-interaction pairs. Next, we chose the split luciferase complementation assay to determine their interactions because of the high dynamic range, low background, commercially available substrates, and compatibility with experiments in living cells^38,39^. This method requires a systematic toggling between the N- and C-terminal fusion and between the fusion of the larger or smaller fragments of the split luciferase to a given protein of interest, because we would not know *a priori* the orientation of the two binding partners, and placing the split luciferase fragments on the wrong termini of the binding partners may sterically hinder the interaction and lead to false negatives. Also, adding a fusion partner may interfere with the native folding of the protein under assay, and thus it is desirable to screen both the large and small fragments for a given protein. For heterologous interaction pairs, this systematic toggling would multiply the number of conditions by 8, and for self-interaction pairs, by 4, which brought the total number of conditions to 484. Thirdly, a critical parameter to assess for GV assembly is the potential interdependence of an interaction pair on a third protein, because this will be the path to identifying protein complexes and constructing a roadmap of the assembly pathway. As the first step to probe the existence of such interdependence relations, we would assess the interaction of a pair of proteins under two conditions: without any background GV proteins and in the presence of all of them. If an interdependence condition exists, we would expect to see the former condition shows a negative result while the latter shows a positive. Adding this step would double the number of conditions, bringing the total number to 968. Lastly, if an interdependence condition is identified, it would be beneficial to probe the interdependence on individual GV proteins, which could add another 9 to 10 conditions to test for each specific interaction pair. We grouped these as “additional” conditions that would be decided case by case.

To cope with this large number of conditions, we decided to leverage the fact that multiple plasmids could be co-transformed into *E. coli*, which allows us to minimize the number of plasmids needed to construct and instead, rely on a mix-and-match of the plasmids to cover all the conditions. Thus, a three-plasmid system was designed and tested, of which each plasmid carried a different antibiotic resistance marker and origin of replication (**Figure 1E and S1**). Plasmid I and II carried individual gas vesicle proteins fused with either Large BiT (Lg) or Small BiT (Sm), which are the two fragments of the split NanoLuc luciferase^38^. Most importantly, Plasmid III contained either all the other GV proteins or an empty backbone, and this plasmid provided a handle to probe whether an interdependence relationship exists for the two proteins in Plasmid I and II. 67 Plasmids III were constructed initially to cover all the single, double, and full deletions for the 968 conditions expected above, and 14 additional triple-deletion Plasmids III were constructed to probe into the subnetworks of plausible functional complexes (see the sections titled The interaction subnetwork of GvpB-GvpF-GvpG and The interaction subnetwork of GvpB-GvpL). Lastly, as a positive control of the split NanoLuc assay, two plasmids were created that carried FKBP12-Sm and FRB-Lg proteins, which will dimerize strongly in the presence of rapamycin^40^. Overall, this strategy successfully reduced the number of cloned plasmids to 127 (**Table S1**).

### Designing a workflow to maximize the information content

The next task is to establish a workflow that can probe multiple parameters and maximize the information collected. Since growing individual cultures and carrying out protein-protein interaction studies will be a lengthy experimental process, we reasoned that adding a couple of analytical steps in our workflow could maximize the utility of each genetic design and cell culture, providing additional knowledge important to future investigation of the assembly mechanism (**Figure 2A**). Three additional parameters, the cellular burden of GV gene expression, the formation of gas vesicles with proteins perturbed by fusion proteins, and confirmation of protein expression, were probed in addition to the protein-protein interaction study (**Figure 2B**). These additional data provided several interesting findings that will be described in the section titled Datasets on cellular burden, protein expression, and tolerance of fusion. Also, protein-protein interactions were measured at two different time points post-induction that would correspond to different cellular states as described in the next section.

**Fig. 2.**
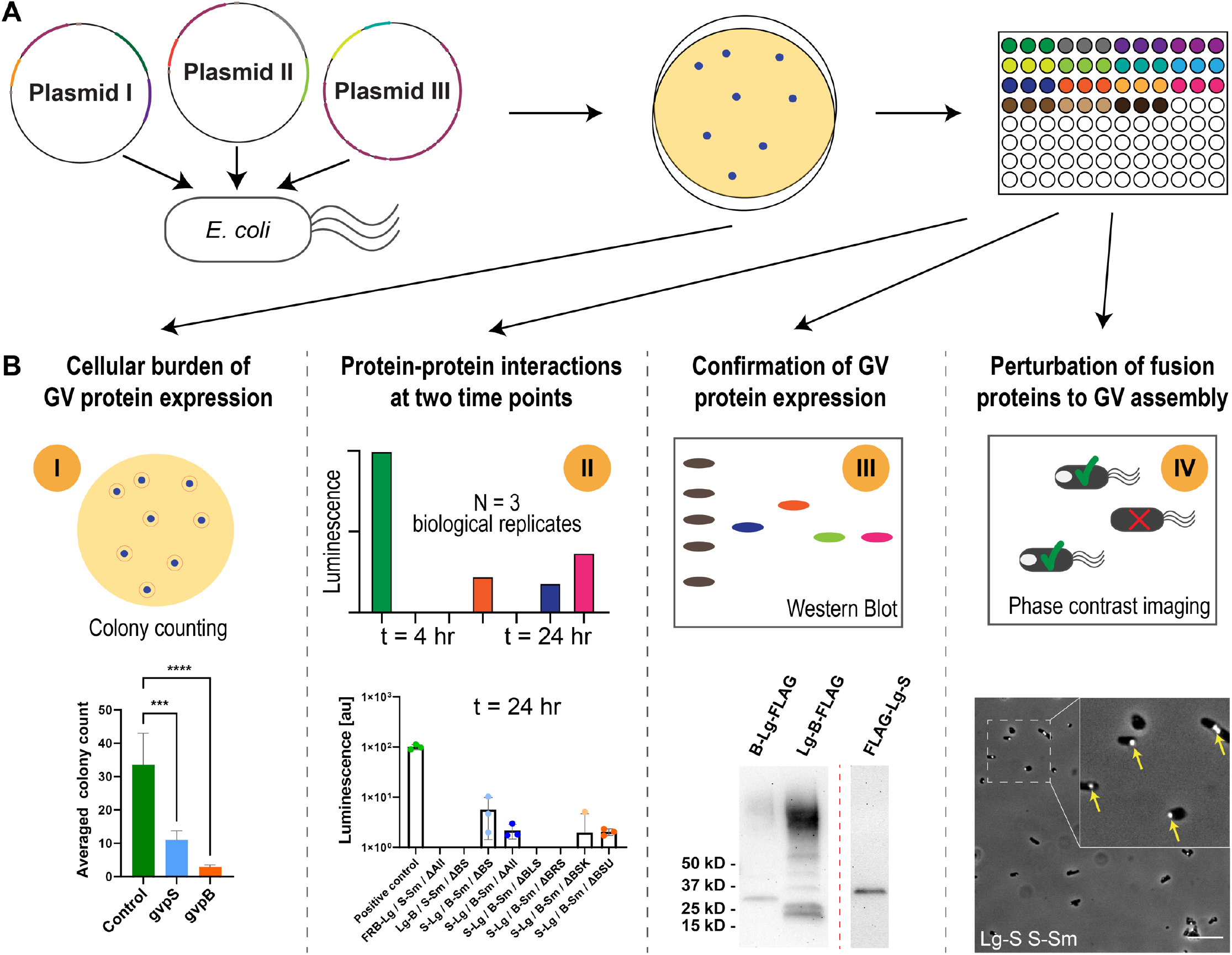
Designing a workflow to maximize the information content. (**A**) Schematic representation of the bacterial culture workflow, including the co-transformation of 3 plasmids (see Fig. 1E) in *E. coli* cells, colony selection on agar plates, and liquid culture in 96-well plates. **(B)** Schematic representation and representative datasets of the assays conducted during and after the bacterial culture. The four major types of data are labeled as I, II, III, and IV in the yellow circles, which include (I) sampling the cellular burden by colony counting on the agar plate, (II) split luciferase complementation assay at two different time points during the cell culturing, (III) Western Blot of selected cell samples to assay the GV protein expression, and (IV) measuring the presence of successfully assembled GVs by phase-contrast imaging as an indication of the tolerance of fusion by the GV assembly factors. Dataset II is used to construct the various interaction roadmaps, while Datasets I, III, and IV are collected to maximize information content and discussed in the section entitled Datasets on cellular burden, protein expression, and tolerance of fusion.

The last action for the experimental design was to minimize the number of protein-protein interaction assays needed. To this end, we prioritized assaying the constructs in which Large BiT was fused with the larger protein of the two interaction partners and Small BiT with the smaller one. We reasoned that the larger fusion protein is usually tolerated better by the larger GV protein. The major shell protein, GvpB, was set as the exception, for which we assayed all configurations, because GvpB is central in the assembly process. For simplicity, the initial testing set included the split NanoLuc all fused to the C-terminus of GV proteins before carrying out additional screening of the fusion at N/C, C/N, or N/N terminus of GV proteins. We reasoned that if a positive interaction is observed under any of the initial C/C fusion conditions, we would no longer need to test the additional fusion constructs.

### Systematic protein-protein interaction maps

With the abovementioned experimental workflow, we ended up testing a total of 1008 protein-protein interactions, each with N = 3 biological replicates. Each experiment included a buffer blank, a positive control that assayed rapamycin-induced FKBP-FRB dimerization, a negative control that assayed the dimerization in the absence of rapamycin, and another negative control of FKBP or FRB fusion protein with a GV fusion protein. To quantify the protein-protein interactions, we defined the rapamycin-inducible positive control in each experiment as 100% strength and scored signals as a fraction of the control. Interactions >20% were defined as strong, 10-20% as significant, and 5-10% as notable. Signals <5% of the positive control were excluded from our final analysis to minimize the chance of including noise-level measurements in our data. In this way, 25 interactions were grouped as “highly relevant”, 76 interactions were grouped as “relevant”, and 43 interactions were grouped as “notable”, resulting in 124 out of 1008, or approximately 10% of all measurements being counted as protein-protein interactions within the operon. Redundant interactions were observed between two GV proteins of different configurations of fusion, and we consolidated these interactions by taking the strongest one as the representative interaction, which resulted in a final set of 14 highly relevant, 8 relevant, and 14 notable interactions. Half of all possible interactions (36 out of 66) of GV proteins have been observed, indicating a dense network of protein-protein interactions (**Figure 3A-C, and Table S2**). Notably, cells were assayed at two different time points in our experimental workflow, and a positive interaction observed in either time point will indicate an interaction between the two GV proteins.

**Fig. 3.**
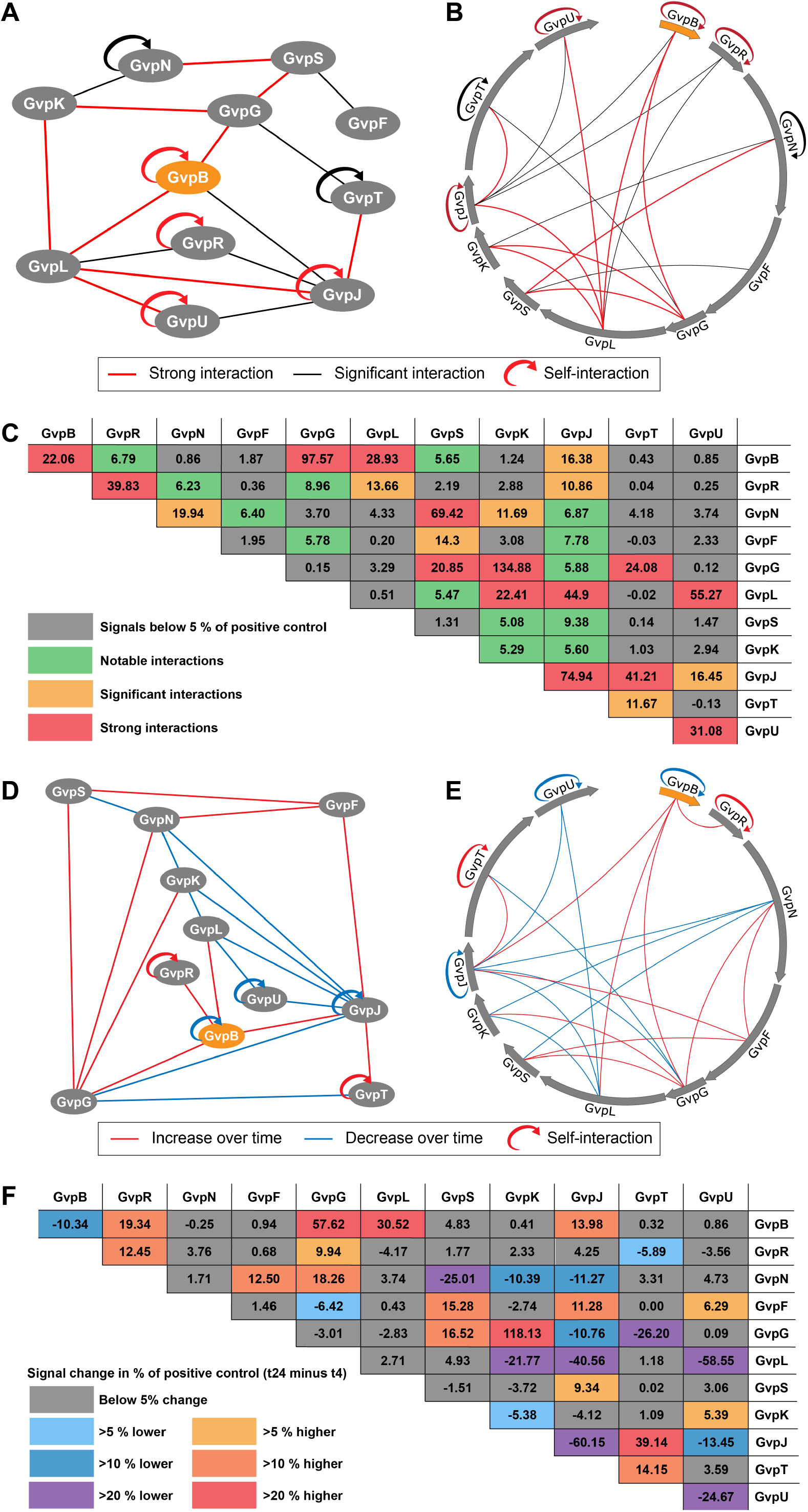
Systematic protein-protein interaction maps. (**A** and **B**) Two styles of representations of the GV protein interaction network. The major shell protein GvpB is represented as an orange oval, and all other GV proteins as gray ovals. In (**B)**, GV proteins are displayed in a circle extracted from the DNA sequence map of the pNL29 GV operon with the actual order and length of the GV proteins preserved. The interactions between proteins are indicated by black lines if they are higher than 10% (significant interaction) of the positive control and red lines if higher than 20% (strong interaction) of the positive control. Half-circular arrows represent the observed self-interactions, which indicate the oligomeric states of the proteins. **(C)** A summary table of the GV protein interactions. For each pair of GV proteins, the highest measured interaction in % of positive control is plotted. Each cell in the table is color-coded to differentiate the 4 categories of interaction strengths as labeled in the legend. **(D** and **E)** Representations of the change of interaction network over time. A red line is used for interactions where the signal is increased by at least 10 % of the positive control (in absolute terms) at t_24_ over t_4_, and a blue line if the signal is decreased by at least 10 %. **(F)** A summary table of the interactions changed over time. Yellow to red colors are used to label an increase in interaction strength, and blue to violet colors for a decrease in interaction strength as labeled in the legend. See also the full list of interaction data in **Table S2**.

Next, we compare the interaction strength at the two different time points, 4 hours (t_4_) and 24 hours (t_24_) post-induction, respectively. We anticipated that t_4_ and t_24_ would reveal different states of protein-protein interactions, because t_4_ corresponds to the initial assembly stage when GVs start to emerge in cells, while t_24_ would correspond to the condition of fully assembled GVs^37^. Furthermore, we hypothesized that the majority of the GV shell proteins might have assembled into GV particles at t_24_, and thus the remaining ones would occupy less of their interaction partners, such as the plausible chaperone protein, which would free up these proteins to engage in other interactions. Among the interactions that showed a strong dependence on time (**Figure 3D-F**), GvpF had an increase in all three interactions over time. In combination with a previous study that suggested direct interaction of GvpF with the truncated α1 helix of GvpA^41^, it is plausible to hypothesize that GvpF functions as a chaperone of GvpB, and more GvpF may have been freed up after the assembly of GvpB into intact GVs. Notably, GvpF was not observed to interact directly with GvpB in our initial t_4_ and t_24_ datasets, and this was investigated further (see section entitled The Interaction Subnetworks of GvpB-F-G).

### Distinguishing direct *versus* indirect interactions of GV proteins

Next, we seek to determine whether an observed interaction is interdependent on a third protein (indirect interaction) or is not influenced by the absence of other proteins, which would suggest a direct interaction. The information can be critical for subsequent studies such as the construction of hierarchical interaction events and structural investigation of the binding sites. Since the roadmaps presented above were constructed in the presence of all 11 GV proteins, we anticipate that only a subset of the interactions were direct ones. Experimentally, we investigated only those pairs that had already shown an interaction in our previous experiments, except for the major shell protein GvpB, of which we screened the interaction with all other GV proteins even if we had not seen an interaction. We also reasoned that keeping the 3 plasmid system will best retain a similar experimental condition for the cells such as the presence of all three antibiotics, and thus we created a decoy Plasmid III that contained no GV protein but only the plasmid backbone (ΔAll, Figure 1). In the data (**Figure 4A-D**), we observed both the cases where the removal of the background GV proteins diminished interactions as expected for the cases of indirect interaction pairs, *e*.*g*., GvpF and GvpN, and the cases where stronger interactions emerged after the removal of the other GV proteins, *e*.*g*., GvpB and GvpF, which were surprising and deserved more investigation (see the next section). Overall, the interactions involving GvpB and GvpJ saw the highest number of increases. Since the major shell protein, GvpB, has a highly hydrophobic N-terminal region that forms the inner surface of GVs^1^, and GvpJ bears substantial homology in this hydrophobic region of GvpB (**Figure 4E**), our data might reveal the fact that in the absence of other GV proteins to chaperone GvpB and GvpJ, their interactions with any given binding partner would become stronger. In contrast, GvpS, which is the only other protein in the operon that has sequence homology to GvpB, showed two strongly reduced interactions together with a couple of weakly increased interactions after the deletion of the background GV proteins. The contrasting behavior of GvpS, GvpJ, and GvpB was further corroborated by the biochemical data that showed cellular burden only from GvpB and a tolerance of GvpS fusion for GV assembly (see the section entitled Datasets on cellular burden, protein expression, and tolerance of fusion). Together, these data suggested different behavior of GvpJ and GvpS compared to GvpB despite the homologous sequence.

**Fig. 4.**
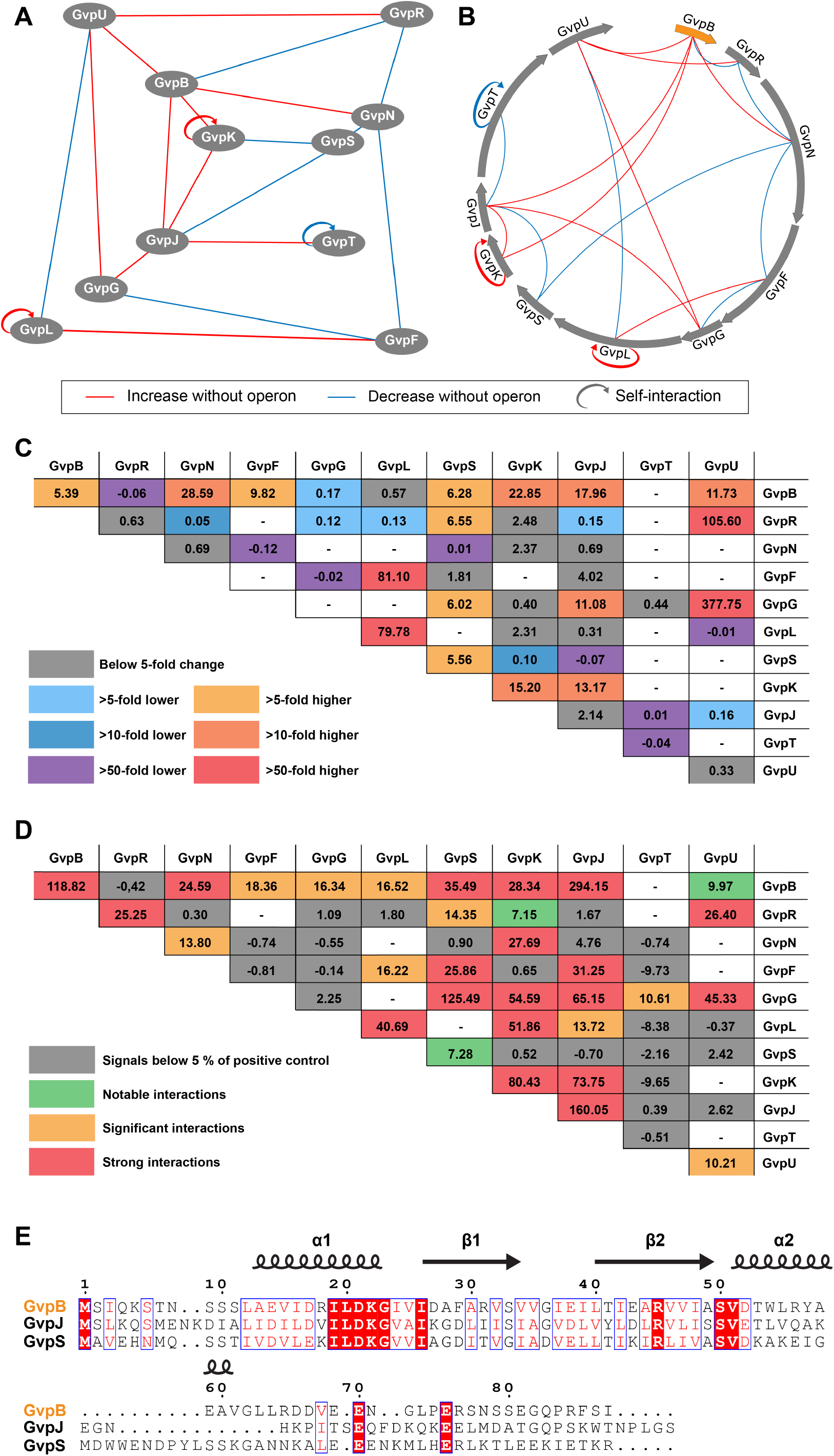
Distinguishing direct *versus* indirect interactions of GV proteins. **(A** and **B)** Two styles of representation of the change of protein-protein interactions when the background GV proteins are removed. Blue lines indicate interactions that are at least 10-fold lower in the absence of background GV proteins, and red lines indicate at least 10-fold higher. **(C)** A summary table of the protein interaction fold changes in the absence of the background GV protein. The fold change is calculated as the signal in the presence of the background GV protein divided by that in the absence. Datapoints are omitted if interaction strength in neither condition is significant, since dividing datapoints of low interaction strength can generate spurious high fold changes. Yellow to red colors are used to label an increase in interaction strength, and blue to violet colors for a decrease in interaction strength as labeled in the legend. **(D)** A summary table of the normalized interaction strength signal in the absence of the background GV protein. Each cell in the table is color-coded analogous to **Figure 3C** to differentiate the 4 categories labeled in the legend. **(E)** Protein sequence alignment of the major shell protein GvpB and two assembly factors, GvpJ and GvpS. The secondary structure of GvpB indicated above the protein sequence was extracted from the cryo-EM structure (PDB: 7R1C)^1^. Alignment was performed using *Clustal Omega*^4^, and the figure was generated using *ESPript*^13^.

### The interaction subnetwork of GvpB-GvpF-GvpG

Next, we proceeded to probe the subnetworks of interactions among the few intriguing clusters of proteins identified from the above roadmaps. Experimentally, we created a set of triple deletion Plasmid III that contained the GV operon excluding the two proteins of interest and a third protein that is hypothesized to mediate their interaction. This strategy allows us to probe the interdependence of certain protein-protein interactions on individual GV proteins. Notably, in our initial interaction screening, we did not observe an interaction between GvpB and GvpF (**Figure 3A-C**), contrary to a previous report that GvpF was the sole interaction partner of the major shell protein^41^. Surprisingly, a strong interaction of GvpB and GvpF re-emerged upon deletion of all other proteins in the GV operon (**Figure 4D**). This hints at the possibility that one or a few of the GV proteins may interfere with the GvpB-GvpF complex, contrary to our assumption that a third protein usually bridges the formation of a complex. To uncover which GV protein is playing this role, we first compared those proteins that interacted with both GvpB and GvpF and found out GvpG had the highest overall interaction (**Figure 3C**). We then created a new plasmid, ΔBFG, of the GV operon with triple deletion of GvpB, GvpF, and GvpG, and indeed, the deletion of GvpG was sufficient to resurrect the GvpB-GvpF interaction (**Figure 5A**), which strongly suggested that GvpF was outcompeted for binding of GvpB in the presence of GvpG. In parallel, another intriguing observation made in the last section was that, while GvpB saw primarily an increase in interactions when other GV proteins were removed, its interaction with GvpG was the only one that decreased (**Figure 4C)**. Thus, we hypothesize that GvpB-GvpF interaction may be a prerequisite to the interaction of GvpB-GvpG. We went on to test the interaction of GvpB and GvpG in the presence of all other GV proteins except for GvpF, and observed that the deletion of GvpF alone was sufficient to attenuate the interaction to a level similar to the ΔAll condition. From these two results, we postulate that GvpF is the primary interaction partner of GvpB, and the formation of the GvpB-GvpF complex recruits GvpG, which in turn supersedes the binding of GvpF (**Figure 5B**).

**Fig. 5.**
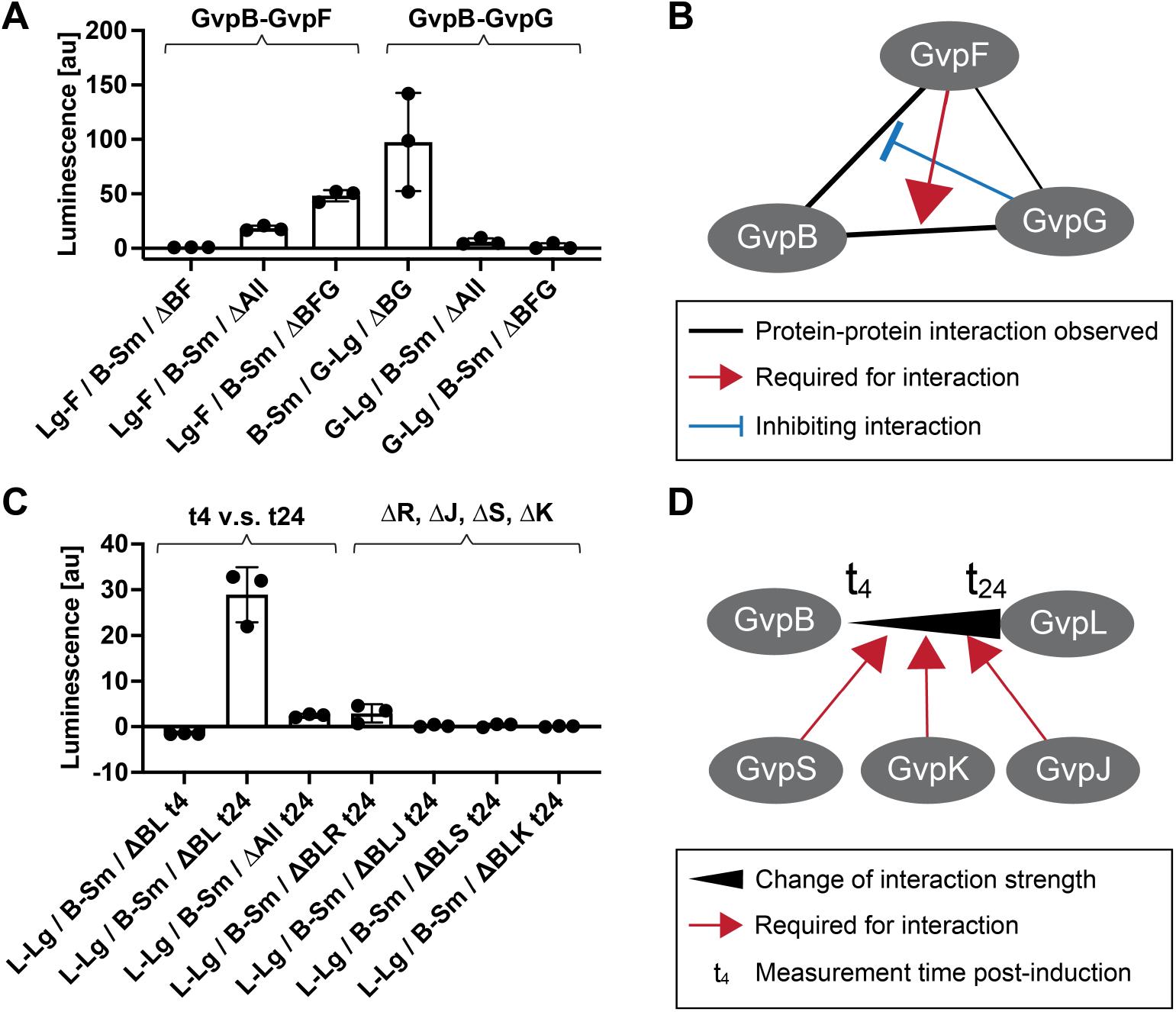
Targeted deletion of GV proteins reveals interaction subnetworks in the GV operon. **(A)** Protein-protein interaction measurements for the subnetwork of GvpB-GvpF-GvpG. The interactions between GvpB fused to Small BiT (B-Sm) and GvpF fused to Large BiT (Lg-F) were measured in the presence of all other GV proteins (ΔBF), in the absence of other GV proteins (ΔAll), and in the presence of all other GV proteins except for GvpG (ΔBFG). Similarly, the interactions between B-Sm and GvpG fused to Large BiT (G-Lg) were measured in the presence of all other GV proteins (ΔBG), in the absence of other GV proteins (ΔAll), and in the presence of all other GV proteins except for GvpF (ΔBFG). **(B)** The postulated interaction map of GvpB-GvpF-GvpG. The types of interactions are labeled in the legend below the map. **(C)** Protein-protein interaction measurements for the subnetwork of GvpB-GvpL. The interactions between B-Sm and GvpL fused to Large BiT (L-Lg) were observed in the presence of all other GV proteins (ΔBL) at 4 hr and 24 hr post-induction (t4 and t24), in the absence of all other GV proteins at 24 hr post-induction (ΔAll t24), and in the presence of all other GV proteins except for single knockout of GvpR (ΔBLR), GvpJ (ΔBLJ), GvpS (ΔBLS), and GvpK (ΔBLK). The interactions are separated into two groups labeled on the top of the graph for the investigation of the change of interaction over time (t4 v.s. t24) and the investigation of the dependence of the interaction on individual GV protein (ΔR, ΔJ, ΔS, ΔK). **(D)** The postulated interaction map of GvpB-GvpL.

It is interesting to compare our observations with a previous study on GV proteins from haloarchaea, which did not include other GV proteins in the background^41^. For the BFG interaction subnetwork, this study observed the interaction of the shell protein GvpA with GvpF, but not with GvpG. The GvpA-GvpF interaction is in agreement with our GvpB-GvpF interaction in the absence of background GV proteins, and the reported absence of the GvpA-GvpG interaction is also consistent with our finding that GvpB and GvpG do not interact in the absence of background GV proteins. However, the presence of the GV operon significantly altered the outcome of the interactions, leading to the discovery of the intriguing interdependent relation among the three proteins. This highlights the importance of modulating background GV proteins in studying the interaction of GV gene clusters.

### The interaction subnetwork of GvpB-GvpL

Next, we investigated the observation that GvpB and GvpL do not interact at timepoint t_4_ but interact strongly at t_24_, as this is one of the pairs that showed a substantial time-dependent change of interactions (**Figure 3F and 5C**). Also, the t_24_ dataset revealed that deleting background GV proteins will substantially attenuate GvpB-GvpL interaction (**Figure 5C**). Thus, we hypothesized that additional GV proteins may mediate the GvpB-GvpL interaction, and this interaction may also be dependent on the stage of GV assembly. Experimentally, we individually removed 4 candidate proteins and created the triple deletion plasmids of ΔBLS, ΔBLK, ΔBLJ, and ΔBLR. Except for GvpR, deleting any one of the other three proteins abolished the GvpB-GvpL interaction (**Figure 5C**), indicating that GvpS, GvpK, and GvpJ all mediate the interaction between GvpB and GvpL. This is further corroborated by the observation of strong interaction of GvpL with GvpS, GvpK and GvpJ especially at the early time point (t_4_) (**Figure 3C and 3F**). Thus, we postulate that GvpB first interacts with GvpS, GvpK, and GvpJ, and the resulting protein complex may be essential for the GvpB-GvpL interaction at a later stage of GV assembly (**Figure 5D**). Notably, among this protein cluster, GvpL was modeled to have a similar structure as GvpF^42^, while GvpB, GvpJ, and GvpS have sequence homology. Thus, based on the previous observation that GvpF interacts with GvpB, it is reasonable to speculate that GvpL may interact with GvpB, GvpJ, and GvpS in a similar fashion. However, this would leave the interesting question of why there are homologous yet essential proteins in the GV operon and what different roles they may play during GV assembly.

### Datasets on cellular burden, protein expression, and tolerance of fusion

First, we hypothesized that the number of *E. coli* transformants may reflect the cellular burden of the GV proteins being expressed, since leaky expression can still occur without chemical induction during the growth of cells on agar plates. Among GV proteins, the major shell protein, GvpB, has a highly hydrophobic surface, which is known to cause aggregation and proteotoxicity^43^. Indeed, we observed that every transformation that included expression of a fusion protein of GvpB led to a decrease in the total number of transformants (**Figure 6A and Table S3**). None of the other GV proteins showed a substantial decrease of transformation efficiency, supporting their roles as assembly factors and minor constituents of GVs. It was somewhat surprising for GvpJ and GvpS, which bear high sequence homology to GvpB and were expected to have similar exposed hydrophobic surfaces. However, our colony counting dataset hinted that biochemically GvpJ and GvpS may behave differently from GvpB and have less exposed hydrophobic surface.

**Fig. 6.**
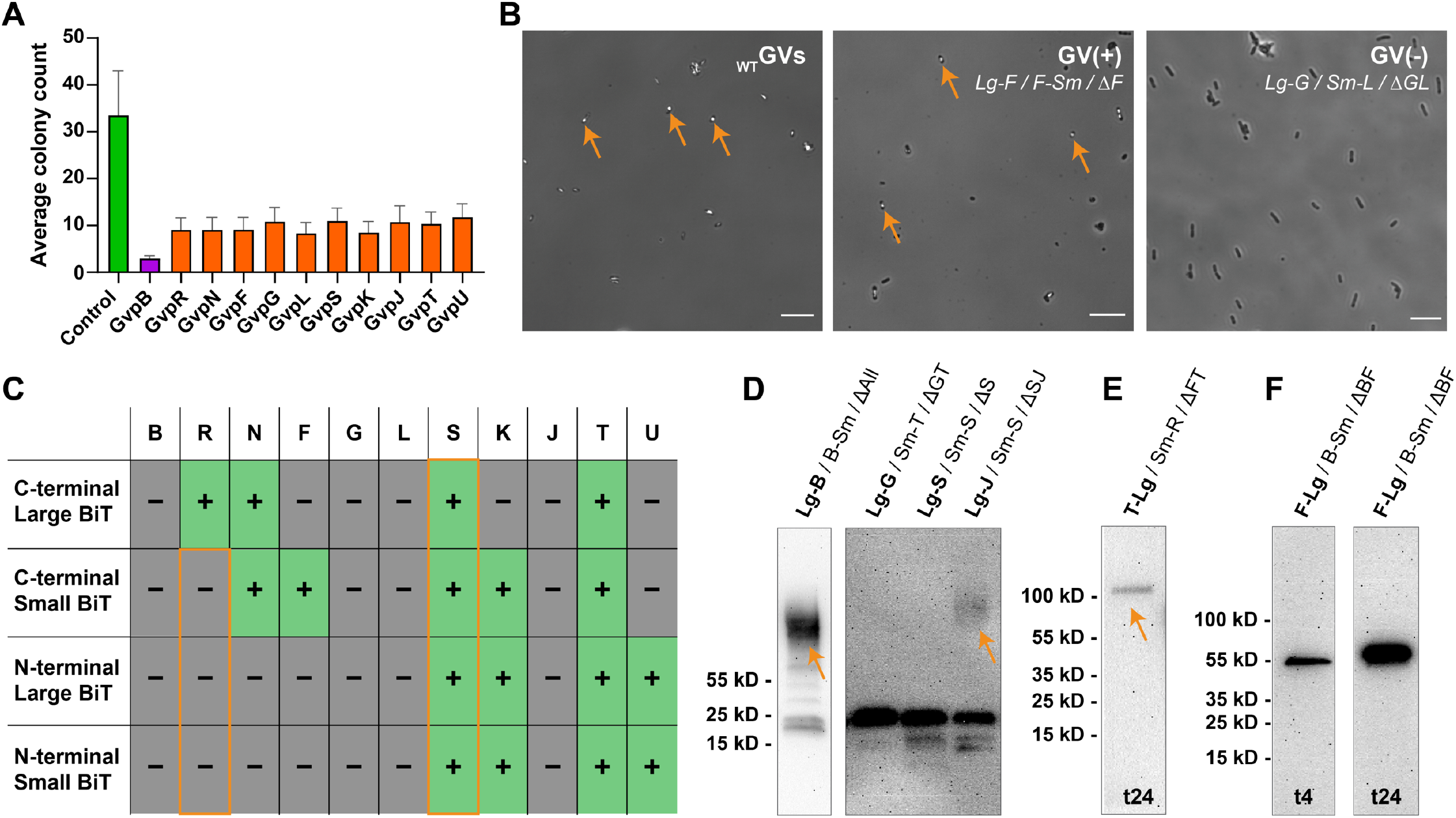
Datasets on cellular burdens, protein expression, and tolerance of fusion. (**A**) The colony counts after transformation indicate cellular burden caused by expressing each GV protein. The colony counts are binned into three groups, and at least three biological replicates and both N- and C-terminal fusion forms are included for each protein. See the full list in **Table S3**. (**B**) Example phase-contrast images of *E. coli*. Orange arrows indicate the representative *E. coli* cells that contain successfully assembled GVs, which are usually revealed as intracellular bright spots. Scale bars represent 20 μm. (**C**) A summary table of whether successfully assembled GVs were observed in phase-contrast images for all 4 types of fusion constructs and all the GV proteins, providing clues for whether a specific fusion will perturb the GV assembly process. See also **Table S3** for the full list of all the conditions tested for phase-contrast imaging and **Figure S3** for representative phase-contrast images of all the conditions containing successfully assembled GVs. (**D**) Example Western blots of the high molecular weight smeared bands of GvpB and GvpJ (labeled by orange arrows). (**E)** Western blot of dimer GvpT-Lg fusion proteins (labeled by an orange arrow). (**F**) Western blots of the GvpF-Lg fusion protein at t4 and t24, representing an example of the increasing protein expression level over time. See also **Figure S2** for additional Western blot images of GV protein -Large BiT fusion used in this study.

Secondly, we assess whether the assembly process of GVs is perturbed by the fusion proteins to individual GV proteins. Making genetic fusions is a common tool to investigate the function, localization, and structure of proteins; however, one could also risk the perturbation of the native function of the protein by the fused partners. For GV proteins, there has not been systematic testing of the tolerance of fusion proteins. Thus, since we have already created all the constructs with both N- and C-terminal fusion of either the Large BiT, a large globular protein, or the Small BiT, a short 11-amino-acid peptide, we decided to take this opportunity to investigate whether these large and small fusion partners will perturb the assembly process of GVs. Experimentally, the successful assembly of GVs can be determined by phase-contrast imaging of bacteria, since the gas compartments of GVs scatter light and produce a distinguishable white spot on these images^44^ (**Figure 6B**). We focused only on conditions where all the GV proteins were provided in the cell and for each configuration of GV fusion proteins, we sampled at least 3 independent cultures at t_24_. A successful assembly should be observed if two conditions were met: (i) the presence of the fusion partner did not interfere with the function of proteins and (ii) the expression level of the fusion proteins does not perturb the stoichiometry of the GV proteins to be beyond the range tolerated by the assembly process. Notably, (i) can also be satisfied if the target GV protein is not essential for GV formation. Since our experiments always have two of the GV proteins simultaneously fused to Large BiT and Small BiT, the observation of assembled GVs will indicate that both fusion proteins were permissive to GV formation, whereas no GVs observed indicated that at least one of the fusion proteins interfered with GV formation. Screening through 149 conditions led to the discovery that 17 conditions permitted the formation of GVs, and grouping these data allowed us to identify which fusion proteins were suitable for GV formation by elimination (**Figure 6C** and **Table S4**). First, we observed that all fusion proteins of GvpS and GvpT permitted GV formation. While GvpT was known to be non-essential to the formation of GVs^6^, it was surprising to see that an essential protein, GvpS, allowed for GV formation to a level close to the wildtype in all the fusion configurations. Moreover, GvpS bears high homology to GvpB on the N-terminal hydrophobic region (**Figure 4E**), and while none of the GvpB fusions gave intact GVs, all GvpS ones permitted the assembly. Another surprising result was that, while GvpR was previously determined to be non-essential^6^, we observed that 3 out of the 4 fusion configurations of GvpR would perturb GV formation. Thus, the creation of less-functional GvpR might cause more disturbance to the assembly process of GVs than simply eliminating GvpR. We anticipate these finding would pave the way to uncover the functional role of GvpR in the assembly process of GVs. For all other proteins, the site of fusion and size of the split-luciferase fragment influenced GV formation as expected, and all together, these experimental results will serve as important guides to the future design of fusion proteins to study the function and cellular location of these proteins.

Lastly, to monitor the expression levels of our fusion protein constructs and to exclude that an absence of luciferase signal is due to poor protein expression, we appended a FLAG-tag on the distal end of each Large BiT to allow the determination of protein expression by Western blot. We observed that all fusion proteins were expressed and appeared at the expected molecular weight with the exception of the major GV shell protein GvpB and its homolog, GvpJ. They were observed as smeared bands at high molecular weight, suggesting that they were insoluble in SDS (**Figure 6D**), and this agrees with previous studies on SDS-PAGE of GV major shell protein^45^. We also observed a dimer form of GvpT (**Figure 6E**), which agrees with the observation of strong self-interaction of GvpT at t_24_. Lastly, we observed a general trend for higher expression levels at t_24_ than t_4_, in line with the higher raw luminescence intensity at t_24_ (**Figure 6F**). Notably, higher expression levels at t_24_ also hold true for the positive control samples, supporting the choice of using the percentile rather than raw luminescence intensity as the quantification method.

## DISCUSSION

In this work, we established a systematic interaction roadmap of GVs that included the interdependence of the protein interactions on the presence of other background assembly factor proteins. Such a protein-protein interaction network is often one of the first steps to understanding the assembly mechanism of a protein organelle. For example, delineating the protein-protein interactions of β-carboxysome biogenesis revealed the core-first assembly pathway^46^, which paved the way to many subsequent studies on the structure and function of individual carboxysome proteins, the redesign of the enzymatic core and shell protein scaffolds, and engineering of carboxysome-like organelles for metabolic engineering^47-49^. Similarly, we anticipate that this work of GV protein interaction roadmap will lay the foundation for future studies and engineering of GVs. To give some specific examples, understanding the protein interactions during GV assembly will guide the optimization of the heterologous expression in therapeutically relevant cells by informing the interdependence of protein players, of which the stoichiometry may be aligned and the binding interface may need to be optimized in a particular cellular context. The poor assembly efficiency during heterologous expression is currently the main obstacle hindering the biotechnological applications of GVs, exemplified by the finding that several assembly factor proteins need to be supplemented in a “booster” plasmid during the mammalian expression of GVs^6^. As another example, this work provided systematic information on the tolerance of GV proteins for terminal fusions, which may be the sites for adding designed proteins to GVs. As the third example, constructing the protein interaction roadmap in this work will be an essential step toward *in vitro* reconstitution of GVs, which will free up the current limitation that GVs have to be manufactured inside a cell. While the current understanding of the biology of GVs is still in its infancy, we anticipate a rapid growth of studies on this topic to match the recent surge of biotechnological applications of GVs.

Building on current knowledge, we would like to postulate an assembly process of GVs encoded in the pNL29 operon (**Figure 7**). **Stage I** would be the initial “seeding” of GVs, which is currently the least understood step in the entire assembly process due to the difficulty of experimentally observing these seeding GV protein complexes. Here we propose that GvpS, GvpK, and GvpJ may be involved in this seeding process of GVs. This is supported by the observation that these three proteins are required for the interaction between GvpB and GvpL (Figure 5D), hinting that they act upstream of the major shell protein GvpB in the generation of new GVs.

**Fig. 7.**
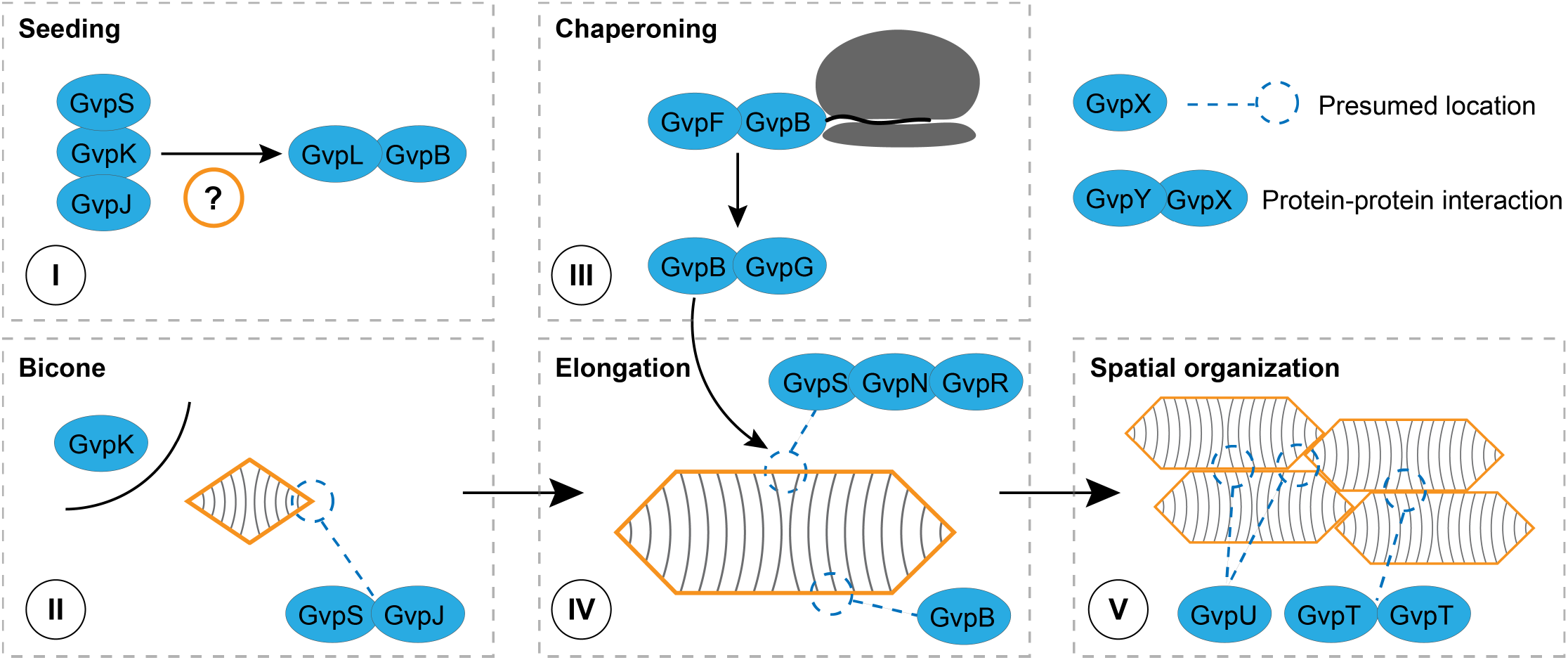
Proposed involvement of GV proteins in each stage of the assembly.

The earliest observable stage of GV growth is the “bicone”^45,50^, which we labeled as **Stage II**, and GvpS, GvpK, and GvpJ likely continue to be involved. Among them, GvpK was notably missing from having a physical attachment to GV particles in a previous mass spectrometry study^51^, suggesting that GvpK is fully soluble in the cytosol, and this corroborates with our observation that most GvpK fusion proteins do not interfere with GV assembly (Figure 6C). Notably, GvpK is one of the few proteins consistently grouped as essential in all 4 major genotypes of GVs studied to date^6,12,52,53^, and yet little information is present to suggest its functional role, and our observation that the presence of GvpK is essential for the GvpB-GvpL interaction (Figure 5D), as well as other notable interactions with GvpN and GvpG, may lead to possible directions to investigate its function. Different from GvpK, GvpS, and GvpJ consist of sequences homologous to the highly hydrophobic, inside-facing α1-β1-β2 segment of GvpB, and thus GvpS and GvpJ may not be fully soluble in the cytosol. While GvpB can fully occupy the middle segment of GVs according to the cryo-EM structures^1,33^, GvpS and GvpJ may occupy the tips of newly formed GVs, and this hypothesis was strongly supported by a recent study on purified bicone GVs, which showed the presence of GvpJ and GvpS in the particles even after washing with 6M urea^26^.

The chaperoning of GvpB, **Stage III**, likely occurs at the ribosome and in parallel to the initiation and elongation process of the GV particles. GvpF is the primary binding partner of GvpB (Figures 4D & 5A), an interaction that is superseded by the secondary binding of GvpG (Figures 3C & 5B). We propose that GvpF and GvpG are the chaperones of GvpB and prevent it from becoming thermodynamically trapped and forming proteotoxic amyloid species^43,54^. GvpF was observed to bind the first α-helix of haloarchaeal shell protein GvpA^41^, supporting a co-translational chaperoning. The presence of GvpG attenuates the binding of GvpF to GvpB (Figures 5A-B), indicating that either they compete for the same binding surface or the fully translated GvpB adopts a conformation with higher affinity to GvpG. It is worth noting that GvpF and GvpG also interact with GvpS and GvpJ and may participate in the Stage (I) and (II) as well. Lastly, GvpF has a homolog, GvpL, and it would be important to investigate why an organism would require 2 copies, or 3 in the case of *Serratia sp*.^55^, of GvpF/L chaperones.

For the elongation of GVs (**Stage IV**), it is established that the insertion of the shell protein GvpB occurs at the middle polarity reversal point observed in the cryo-EM structures^1,33^, and yet how the assembly factor proteins facilitate the insertion process remains unclear. GvpN likely participates, since it is established that ΔGvpN mutants allow the formation of bicone GVs but these GVs do not elongate in the absence of GvpN^26,52,55,56^. GvpN is an AAA+ ATPase, and we observed its self-interaction (Figure 3C), which is in agreement with AAA+ protein-specific homooligomers^57^. AAA+ ATPases are known to promote degradation or refolding of their protein substrates^57^, and we hypothesize that GvpN uses energy to prepare the correct folding of GvpB for insertion into the growing cylindrical portion of GVs, or alternatively, uses energy to proofread the folding of GvpB along GV particles^58^. Since GvpB is the main component of GVs, our observation that GvpN interacts with GvpB and not GvpJ or GvpS in the absence of all other assembly factors (Figure 4C) further support the role of GvpN in the elongation process of GVs. Interestingly, in the presence of all GV proteins, GvpS became the strongest interaction partner of GvpN (Figure 3C), and thus GvpS might also participate in the growth of GVs at the polarity reversal point. Lastly, this interaction of GvpN and GvpS is partially dependent on the presence of GvpR (Table S2), indicating that GvpR may assist the activity of GvpN. The relation between GvpR and GvpN was corroborated by previous reports that GvpO, a haloarchael homolog of GvpR, will influence the expression of GvpA, GvpC, and GvpN^51,59^. However, whether GvpR/O is essential for the assembly process remains controversial and may depend on cellular context: GvpO was grouped as an essential protein in haloarchaea^52^, but for pNL29-encoded GVs, ΔGvpR mutant can produce GVs^6^; and in this work, we additionally observed that most fusion proteins of GvpR would produce GV-negative cells (Figure 6C) indicating an unproductive gain-of-function.

Finally, **Stage V** is the spatial organization of GVs. While this stage is not essential for the formation of GV particles, the spatial organization would be important to minimize GVs’ occupancy of cytosolic space and was recently shown to modulate cellular fitness^60^. GvpU and, to a lesser extent, GvpT mediates the clustering of GVs^60^, and both of them are non-essential proteins in the pNL29 operon^6^. Intriguingly, we still observe interactions between GvpU, L, and J as well as GvpT, G, and J, which link them to the initiation and chaperoning stages of GV formation. One plausible explanation is that the clustering of GVs occurs concurrently with the assembly of GVs, and some of the assembly factor proteins may have even evolved affinity to GvpU and GvpT, which will help their spatial recruitment to the sites of GV assembly in the cytosol. Finally, we observed other notable interactions, including a strong interaction between GvpG and GvpK (Figure 3C) which is unaffected by deleting the rest of the GV operon (Figure 4C). Following the abovementioned binding partners of GvpT and GvpU, we furthermore observed that in the absence of all other GV proteins, GvpU shows an increase in binding to GvpR and GvpG (Figure 4C). Together, these findings indicate that the stages of GV initiation, growth, chaperoning, and spatial organization are not discrete, but may occur in parallel and be organized by crosstalk between proteins. The resolution of the assembly mechanism of GVs will not only be intriguing but also mark a highly significant stride for this class of microbial organelles with remarkable physical properties.

## ACKNOWLEDGMENTS

We would like to thank Thyer lab for providing template plasmids encoding the Spectinomycin resistance gene and the p15A and CloDF13 origins of replication. We thank the Shared Equipment Authority (SEA) at Rice University for the access to core facilities and instruments. This work was supported by the Cancer Prevention and Research Institute of Texas (CPRIT), the NIH (R00 EB024600 and R21 EB033607), the Welch Foundation, G. Harold and Leila Y. Mathers Foundation, and John S. Dunn Foundation. M.I. acknowledges support from German Research Foundation (DFG) Postdoctoral Fellowship.

## AUTHOR CONTRIBUTIONS

Conceptualization, G.J.L., M.I., A.P.A.; Methodology, G.J.L., M.I., A.P.A.; Investigation, M.I., A.P.A., V.T.W, E.D.A., A.H.; Formal Analysis, M.I., A.P.A.; Writing – Original Draft, G.J.L., M.I., A.P.A.; Supervision and Funding Acquisition, G.J.L..

## METHOD DETAILS

### Plasmid construction

DNA plasmids used in this study were generated by Gibson assembly and site-directed mutagenesis utilizing Q5 Hot Start High-Fidelity enzymes, HiFi DNA Assembly Master Mix, and KLD Enzyme Mix (New England Biolabs (NEB), Ipswich, MA). Plasmids were generated based on the pET-26b(+) backbone to contain the large or small fragments of the NanoBiT luciferase (Large BiT or Small BiT) used for the split luciferase complementation assay (Promega, Madison, WI, USA)^38^. Large BiT or Small BiT were fused at the N-or C-terminus with GV proteins, FRB, or FKBP12, the latter two of which were used as a rapamycin-inducible positive control of dimerization^40^. An 18-amino acid linker VSQGSSGGGGSGGGGSSG was used between the two fusion partners for all the constructs^61^. For each construct carrying the Small BiT, the origin of replication was replaced with the p15A origin. For those constructs carrying the Large BiT, the origin of replication was replaced with the CloDF13 origin, and the kanamycin resistance gene was changed to a spectinomycin resistance gene. In parallel, deletion mutants of each GV protein in the pNL29 operon and combinations of two or three thereof were generated as indicated. The pNL29 operon was first cloned from the pST39-pNL29 plasmid (Addgene ID 91696), and the ORF of GvpK overlapping the ORF of GvpS was resolved by moving the start codon of GvpK to the downstream of GvpS, generating a new plasmid named as Mega’ (**Figure S1A**). Subsequently, single, double, triple, and full knock-out variants of the GV operon were generated from the Mega’ plasmid. All the plasmids are listed in **Table S1**, and the DNA sequences of all the parts are listed in **Table S5**. Primer design as well as graphical presentation of plasmid constructs was carried out using SnapGene (GSL Biotech LLC, San Diego, CA, USA).

### Transformation

For molecular cloning, bacterial transformations were carried out using NEB Turbo competent cells, and the transformed cells were isolated on agar plates with the respective antibiotics (50 μg/mL Kanamycin or 75 μg/mL Spectinomycin or 100 μg/mL Carbenicillin). For protein-protein interaction experiments, home-made competent BL21(DE3) cells were generated by Mix & Go *E. coli* Transformation Kit (Zymo Research). For each transformation, 20 μL of competent cells were mixed with approximately 50 ng of each of the 3 plasmids as indicated. Cells were incubated for 30 minutes on ice before heat shock for 30 seconds at 42 °C in a water bath. Cells were left to recover on ice for 5 minutes before the addition of 200 μL SOC media in a 1.5 mL tube, followed by gentle shaking for 4 hours to rescue. The transformation mixture was plated on bacterial agar plates supplemented with 0.5-fold of the previously indicated antibiotics and 1 % (w/v) glucose to grow overnight at 37 °C. Plates with successful transformants were sealed and transferred to 4 °C until further use. For transformants that failed to recover on agar plates, the transformation procedure was repeated, and instead of plating, the resulting mixtures were transferred to 5 mL LB liquid media with the same additives to grow overnight at 30 °C.

### Split luciferase complementation assay

Cell cultures, luminescence measurements, and optical density at 600 nm (OD_600_) measurements were all carried out in 96-well microplates to maximize the throughput. First, *E. coli* BL21(DE3) cultures transformed with the 3 indicated plasmids were inoculated from agar plates or liquid media to 1 mL of liquid media containing 0.5-fold of the three antibiotics and 1% (w/v) glucose. Three separate cultures were raised for each combination to ensure 3 biological replicates for the subsequent experiments. The 1 mL cultures were grown overnight at 30 °C in a sealed 96-deep-well storage plate (SSI Bio Inc.) shaking at 600 rpm. For each experiment, a triplicate of Luria Broth (LB) media without bacterial cells was included as negative control and used subsequently for luminescence and OD_600_ measurement. On the following day, 10 μL of these overnight pre-culture cells were transferred to 190 μL of LB media containing antibiotics and 0.2% glucose in a clear 96-well microplate (#3370, Corning, Corning, NY, USA) and left to grow at 30 °C and 600 rpm until all cultures reached OD_600_ above 0.3 measured in a Biotek Synergy H4 Multimode (Agilent, Santa Clara, CA, USA) microplate reader. Isopropyl b-D-1-thiogalactopyranoside (IPTG) was added to a final concentration of 20 μM in all wells, and this was counted as time zero (t_0_). At 3.5 hours after induction, rapamycin (Biovision, Milpitas, CA, USA) in DMSO was added to a final concentration of 25 μM in wells containing cells with the FRB-Lg / FKBP12-Sm / ΔAll constructs as positive control. At 4 hours and 24 hours after induction, respectively (t_4_ and t_24_), 10 μL of assay cells were mixed with 10 μL of NanoGlo Live Cell Substrate (Promega, Madison, Wisconsin, USA) and 30 μL of LB to make up a total of 50 μL reaction mixture in an opaque, 96-well half-area microplate (#3694, Corning, Corning, NY, USA) for the luminescence measurement. Shortly before the luminescence reading, OD_600_ of all samples was measured in the clear 96-well plate, and samples for SDS-PAGE and phase contrast microscopy were collected as indicated.

### Data processing and evaluation

Data from the split luciferase complementation assay are reported as follows. OD_600_ and luminescence of the blank buffer control were treated as the baseline, and the values are subtracted from all other measurements. Next, for each well, the luminescence measurement was divided by the OD_600_ measurement to normalize the signal for variations in cell density. For each microplate, a positive control was included that contained cells expressing FRB-Lg / FKBP12-Sm / ΔAll plasmids and was supplemented with rapamycin 30 minutes before measurement. The signal from positive control was set to be 100%. Each microplate also contained two negative controls. The two negative controls consisted of cells expressing FRB-Lg with an arbitrary GV protein fused with Small BiT or FKBP12-Sm with an arbitrary GV protein fused with Large BiT. These negative controls are made assuming that GV proteins do not interact with FKBP12 or FRB protein. All measurements lower than the negative control were considered to represent non-interaction, and all measurements of the samples were reported as percentages of the positive control. We noted that some interactions measured were stronger than the positive control, giving rise to values >100%.

### Phase-contrast imaging

For phase-contrast microscopy, 5 μL of the bacterial cultures at t_24_ were transferred to object slides and mixed with a drop of Fluoroshield mounting medium (Sigma-Aldrich). Coverslips were added and samples were left to dry for at least 30 minutes before analysis on an Eclipse TI2 inverted microscope (Nikon, Melville, NY, USA) to identify cells containing GVs. For imaging, the 40x phase-contrast objective and 1.5x tube lens are combined to result in a 60-fold total magnification. The Ph2 condenser annulus was used, and the exposure was held at 700 milliseconds for all the images. Phase-contrast images were prepared for publication with *Fiji*^62^.

### Western blot

30 μL of the indicated bacterial cultures were mixed with 30 μL of 2x Laemmli buffer and incubated at 98 °C for 5 minutes. Samples (using 25 μL each) including 5 μL PageRuler Plus (Thermo Fisher Scientific, Waltham, MA, USA) or Precision Plus Dual Xtra (Bio-Rad, Hercules, CA, USA) prestained protein ladder were applied to SDS-PAGE using 4-20% Mini-PROTEAN TGX precast protein gels (Bio-Rad, Hercules, CA, USA). Western blots were carried out in a Trans-Blot Turbo system using Trans-Blot Turbo Mini 0.2 μm Transfer-Packs (Bio-Rad, Hercules, CA, USA) using the Standard setting on the instrument. Total protein staining was performed using Revert 700 Total Protein Stain (LI-COR, Lincoln, NE; USA) following the manufacturer’s instructions and detected using a FluorChem M imager (Protein Simple, San Jose, CA, USA) by the Multifluor Red setting. Immunodetection was carried out by blocking PVDF membranes in 5% (w/v) fat-free dried milk powder followed by incubation overnight in rat anti-FLAG antibodies diluted 1:500 (Agilent Technologies, Santa Clara, CA, USA). After a brief rinse in Tris-buffered saline with Tween™ 20 detergent (TBST), goat anti-rat HRP-conjugated secondary antibodies (Invitrogen, Waltham, MA, USA) were used in 1:5000 dilution to incubate membranes for 1 hour. Membranes were washed 3x in TBST, followed by TBS, before detection using Pierce ECL Western Blotting Substrate (Thermo Fisher Scientific, Waltham, MA, USA) with the FluorChem M imager using the “Chemi + Markers” setting and exposing for no more than 10 minutes. Western blot images were contrast optimized with *Fiji*^62^.

### Quantification and statistical analysis

The statistical information of the experiments, including sample size (n) and P-values, can be located in the figures and figure captions. The statistical analysis was carried out using GraphPad Prism software and is represented as the mean ± standard deviation (STD).

