## supplemental materials for "Elucidating the Assembly of Gas Vesicles by Systematic Protein-Protein Interaction Analysis"

**Table S1. List of plasmid constructs used in this study.**

The table lists the names, antibiotic resistance genes, origins of replication, and brief descriptions of all plasmid constructs used in this study.

**Table S2. List of all protein-protein interaction data.**

All protein-protein interaction data obtained in this study are shown as tables in 6 Excel workbooks. Workbook 1 lists all interactions measured in the order of how the proteins are present in the pNL29 operon (GvpB, GvpR, GvpN, GvpF, GvpG, GvpL, GvpS, GvpK, GvpJ, GvpT, and GvpU) and sorted by the configuration of the fusion protein. Gray boxes indicate interaction geometries of fusion proteins that were not measured, and the color coding of all interactions measured follows the pattern of Figure 3C. Workbook 2 lists the measurements taken in the absence of the background GV proteins, and the data are organized in the same way as Workbook 1. Workbook 3 provides a direct comparison of the data presented in Workbooks 1 and 2. Workbook 4 lists all interactions measured for triple deletions corresponding to Figure 5 and the sections on the interaction subnetworks. In addition to the two GV proteins being investigated, a third GV protein was deleted from Plasmid III to probe the interdependence of the interaction on this specific GV protein. This third GV protein is labeled in row 1. Workbook 5 is a condensed summary of all the interaction values extracted from the study. Column 1 labels the interaction pair under investigation, and Row 1 labels the conditions under which the interaction is studied. Workbook 6 contains all the data organized in the same style as Workbooks 1-4.

**Table S3. List of data on colony counting.**

Table lists the three plasmids utilized for transformation for all bacterial agar plates on which colony counting was performed. The counts were binned as no colonies, more than 1 colony (labeled “1”), more than 10 colonies (“10”), and more than 100 colonies (“100”). The counts are used as an approximation of the proteotoxic burden of the protein cargo expressed from the transformed plasmids.

**Table S4. Summary of phase-contrast microscopy screening.**

The table lists the three plasmids on which the phase-contrast imaging was performed to evaluate the existence of intact, fully assembled GVs. The phase contrast results were binned as follows: GVs observed in at least one cell (“+”), GVs observed in a significant fraction of cells (“++”), and GVs not observed in any cell (“-“). All those conditions with GVs were grouped at the top of the table.

**Table S5. Genetic sequences of main elements used in the project.**

Table lists the DNA sequences of the parts and proteins used in this study. Column C lists the source from which the DNA sequence was derived, and the relevant previous studies including pST39-pNL29 plasmid[1,2], FKBP12 and FRB[3], Small BiT and Large BiT[4], FLAG tag[5], and the linker sequence[6].

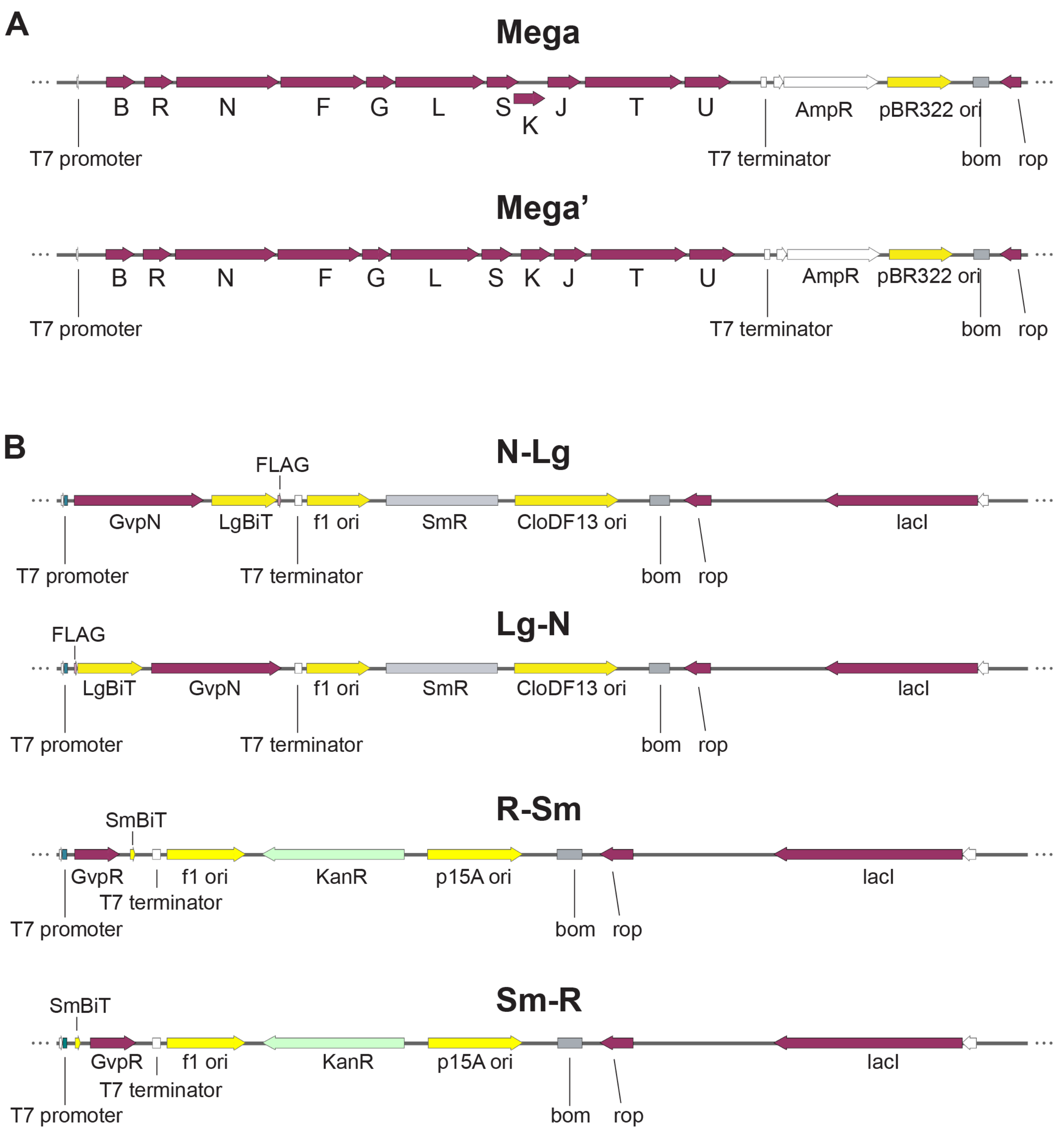


**Figure S1. Overview of plasmid designs used in this study**. Representative plasmid designs are shown in a linear form for ease of view. **(A)** The plasmid Mega encodes the proteins GvpB, GvpR, GvpN, GvpF, GvpG, GvpL, GvpS, GvpK, GvpJ, GvpT, and GvpU (indicated by single letters) which can be expressed to generate recombinant GVs in *E. coli*. The plasmid contains a resistance marker for ampicillin (AmpR) and a pBR322 origin of replication. Mega’ is identical to Mega, but the overlap between GvpS and GvpK has been resolved by molecular cloning. All knockouts of one, two, or three GV proteins used in this study were derived from Mega’. **(B)** The plasmid N-Lg encodes GvpN-Linker-NanoLuc Large BiT-FLAG tag inducible by IPTG. The plasmid contains a resistance marker for Spectinomycin (SmR) and the CloDF13 origin of replication which is not in the same compatibility group as the pBR322 origin. Lg-N is identical to N-Lg but encodes FLAG tag-NanoLuc Large BiT-Linker-GvpN and all other plasmids X-Lg or Lg-X are constructed analogously. The plasmid R-Sm encodes GvpR-Linker-NanoLuc Small BiT inducible by IPTG, and the FLAG tag was not included. The plasmid contains a resistance gene for Kanamycin (KanR) and the p15A origin of replication which is not in the same compatibility group as the pBR322 and CloDF13 origins. Sm-R is identical to R-Sm but encodes NanoLuc Small BiT-Linker-GvpR and all other plasmids X-Sm or Sm-X are constructed analogously.

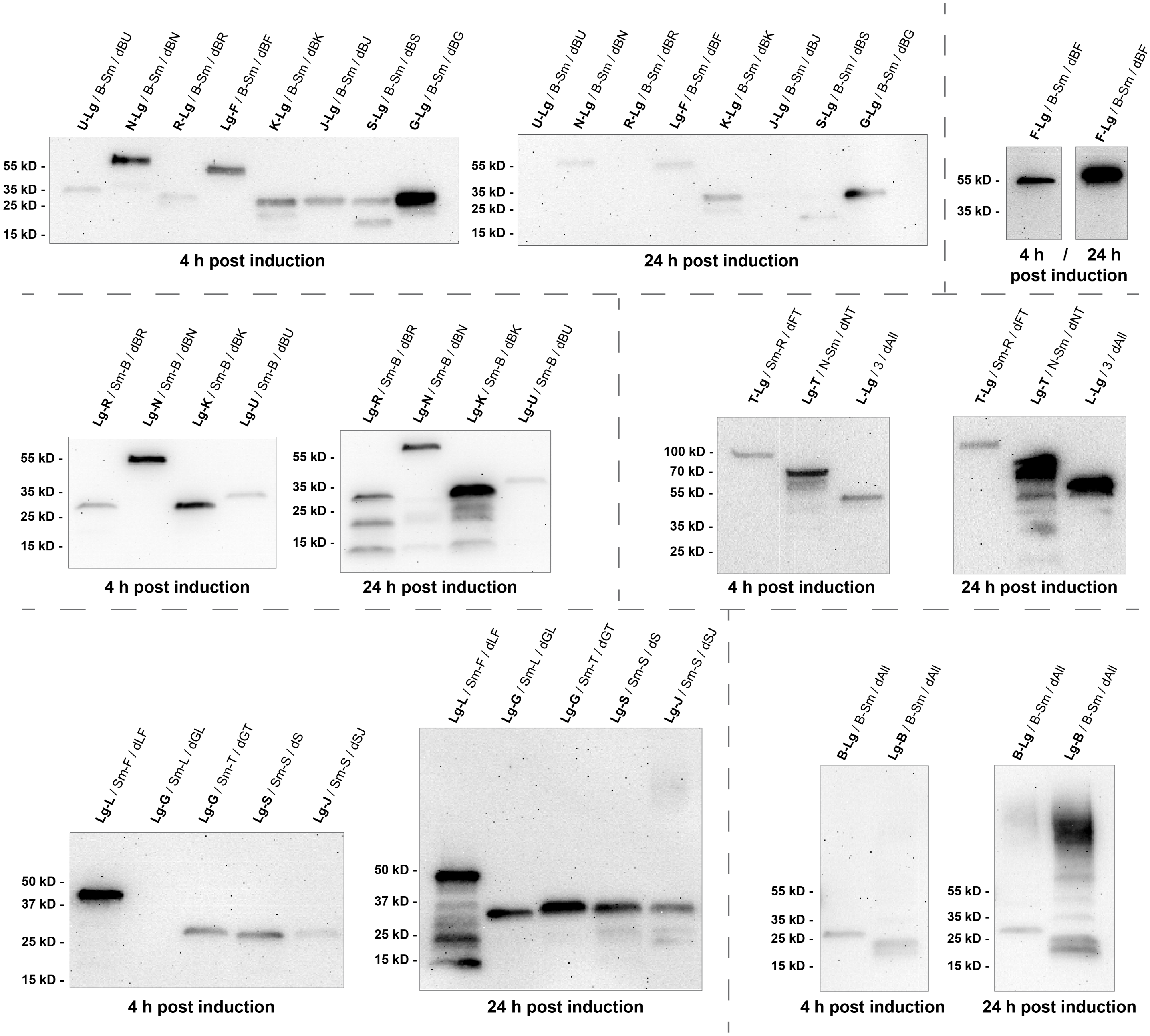


**Figure S2. Western blot data to confirm expression of Gvp fusion proteins.** FLAG tags were always included at the distal end of the Large BiT to allow for the detection of the expression of the fusion protein by western blotting (Figure S1B). We sampled a subset of the samples to confirm the correct expression of our fusion proteins and to exclude the possibility that an absence of signal is due to an absence of protein expression. Each GvpX-Lg and Lg-GvpX was sampled in at least three independent instances and at least one instance of each is sampled for both $t_4$ and $t_{24}$, and the figure shows a representative example out of three for each fusion protein analyzed in this way.

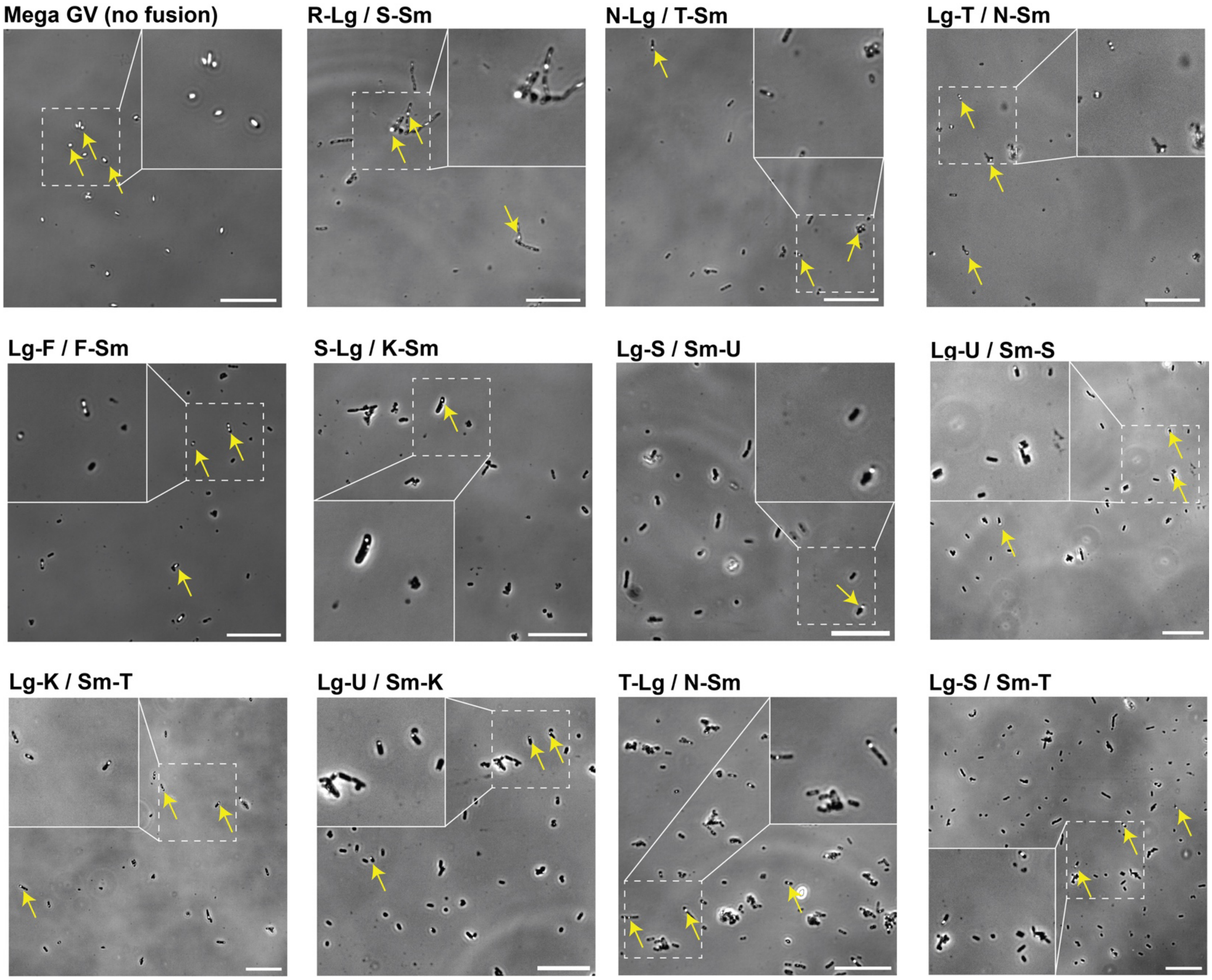


**Figure S3. Phase-contrast images to examine the tolerance of the fusion partners to GV assembly factor proteins.** For each fusion protein, at least three separate samples were taken for phase-contrast microscopy and analyzed for the existence of intact, successfully assembled GVs. We show the expression of GVs from the unmodified pNL29 operon and at least one for each GV fusion protein that gave rise to successfully assembled GVs (Figure 6C). The scale bars represent 20 μm, and the white boxes mark a zoom-in view of the select regions. Arrows indicate the representative GV-containing cells.
